# Temporal ordering of omics and multiomic events inferred from time series data

**DOI:** 10.1101/2020.04.14.040527

**Authors:** Sandeep Kaur, Timothy J. Peters, Pengyi Yang, Laurence Don Wai Luu, Jenny Vuong, James R. Krycer, Seán I. O’Donoghue

## Abstract

Temporal changes in omics events can now be routinely measured, however current analysis methods are often inadequate, especially for multiomics experiments. We report a novel analysis method that can infer event ordering at better temporal resolution than the experiment, and integrates omic events into two concise visualizations (event maps and sparklines). Testing our method gave results well-correlated with prior knowledge and indicated it streamlines analysis of time-series data.

---

A range of emerging omics and multiomics techniques now provide unprecedented ability to systematically track the time course of changes in abundance, for example, in mRNAs, proteins, or posttranslational modifications (PTMs). The resulting time-series data can then be used to infer the ordering of underlying events (e.g., activation of kinases, transcription factors, or translation factors) that explain the observed changes in abundance. These ordered events provide insight into the causal flow of information underlying transitions in cellular state. These insights, in turn, have the potential to lead to fundamental advances in biology and biomedical sciences – however, realizing this potential requires improving the analysis methods used, which are often inadequate with existing data sets^1,2^, especially from multiomics experiments.

When analysing data from such experiments, a common first step is to partition the (typically) tens of thousands of abundance time profiles found into tens of clusters, each with similar temporal changes that are then visualized in a single profile plot^1,2^ (Fig. 1). A wide variety of clustering methods are available, ranging from widely-used, general purpose methods (e.g., fuzzy c-means; FCM^3^) to specialized methods tailored for specific, omics experimental scenarios^4,5,6.^ The specialist methods typically combine clustering with assigning the underlying events, for example, to specific kinases or transcription factors. Often, these events are further assigned to specific time intervals used in the experiment, or to approximate time windows (e.g., early, intermediate, or late response), thus defining an implicit event ordering. However, a key limitation of current approaches is seen when plotting the inferred events using temporal-based layouts^7^: there can often be many more events than time intervals in the original experiment, resulting in many inferred events clustered together at the same time interval. This limitation becomes more pronounced with technologies, such as single-cell RNA sequencing (scRNA-seq) and multiomics experiments^8^, that can result in large numbers of clusters and events. To address this limitation, we have developed Minardo-Model, an automated, reproducible method that uses statistics to infer events ordering – in many cases, achieving better temporal resolution than the experiment. Minardo-Model also uses two novel, compact, and scalable visual representations to create integrated overviews showing all events inferred from a data set. We named our method ‘Minardo-Model’ as it was initially created to help automate generation of cell-based, temporal layouts known as Minardo graphs^1,2,7^, which, in turn, were inspired by Charles Minard’s historic chart of Napoleon’s Russian campaign^1,2^.

**Fig. 1.**
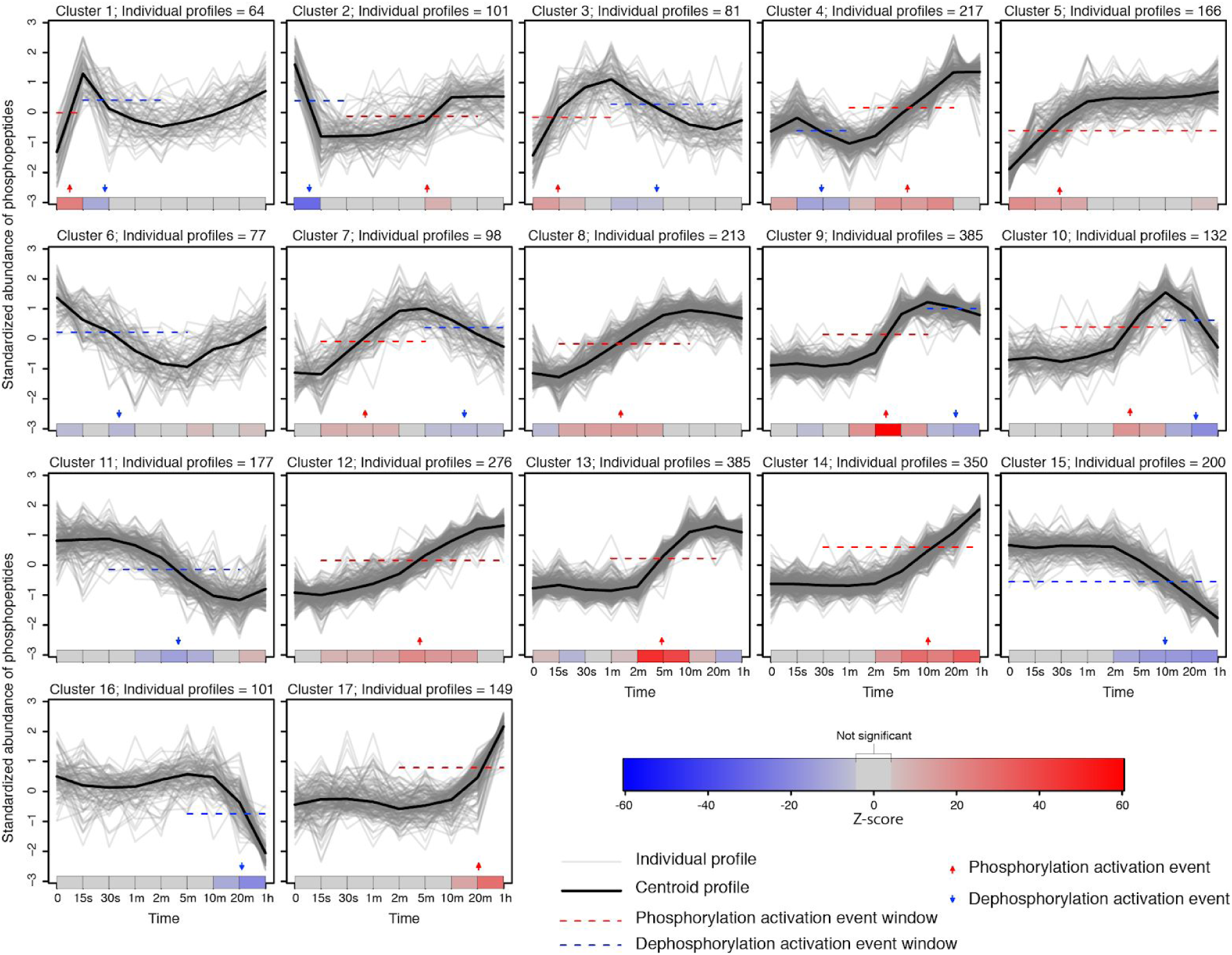
Profile plots of phosphorylation changes in adipocytes following insulin stimulation. Individual profiles for 3,172 phospho-sites grouped into 17 clusters using FCM^3^. Below each cluster, a single-row heat map indicates significant changes in mean phosphopeptide abundance between consecutive time points (red and blue showing phosphorylation and dephosphorylation, respectively), calculated via generalized linear models derived from individual profiles, and using Z-scores from a post-hoc Tukey test. The Z-scores and p-values are used to compute event windows, within which events are defined. Thus, the behavior of each cluster is summarized as a series of discrete phosphorylation and dephosphorylation events (red and blue arrows, respectively), based on the median time at which all individual profiles in a cluster cross half-maximal abundance within each event window (identified by the red or blue dashed line, respectively) – see Supplementary Fig. 3 for details. The ordering of these events was determined statistically, then used to sort and number the clusters. The cluster centroids are shown only to provide a graphical indication of the trend within each cluster. Figures made using data from Humphrey *et al*.^*14*^ with Minardo-Model and edited in Adobe Illustrator.

Minardo-Model takes, as input, time series data that are either already clustered using specialist methods^5,6^, or, if not, the input data are clustered using a generic method (FCM^3^). The following three steps are then applied (see Methods and Supplementary Fig. 1 for details):

1. Generalized linear models (GLMs)^9^ and Tukey post-hoc contrasts^10^ are used to identify a series of event windows within each cluster. These are windows of time during which significant changes in mean abundance occur consecutively in one direction only (either increasing or decreasing), i.e., each window denotes the inferred time span of an underlying event, such as the activation of transcription (Fig. 1 & Supplementary Figs. 4-6).
2. Within each event window, an individual event time is then calculated using each individual profile, defined as the time at which a profile crosses halfway between the lowest and highest median abundance values within each event window (hereafter referred to as the 50% threshold). The distribution of individual event times are shown via density plots (Supplementary Figs. 3, 8-10), which are in turn used to define either the mean or median event time (for normal or skewed distributions, respectively). Finally, the mean or median event time is then indicated as an arrow on the profile plots (Fig. 1 & Supplementary Figs 4-6).
3. False discovery-rate corrected Student t-tests^11^ or Mann-Whitney U comparisons^12^ (for normal or skewed distributions, respectively) are then applied to all pairs of events found in the data set, in each case, testing whether any observed differences are statistically significant. These pairwise comparisons are then analyzed using a graph-traversal strategy, resulting in a unique ordering for all events. These results are plotted using event maps, a novel layout that provides an easy-to-read, visual overview of all inferred clusters and events (Fig. 2a). The results are also plotted using event sparklines, a second novel layout that provides a compact summary showing the ordering of all inferred events within a few lines of text, i.e., at so-called typographic resolution^13^ (Fig. 2b).

We tested Minardo-Model by applying it to two cutting-edge, experimental data sets, described briefly below (see Methods and Supplementary Information for details). Both sets have quite high levels of noisy data, and cover well-characterized processes in which the inferred event ordering could be evaluated against multiple prior publications.

**Fig. 2.**
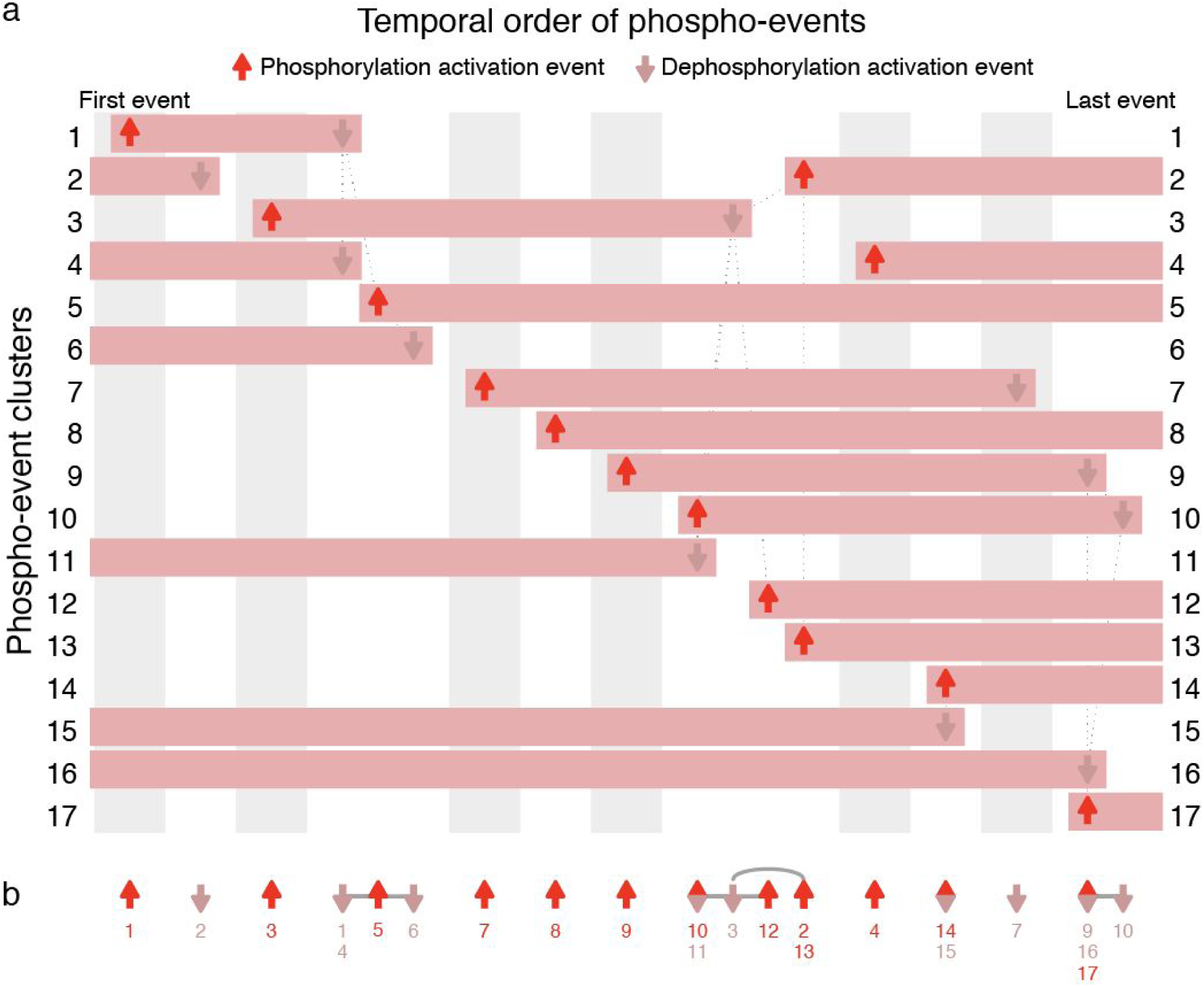
Event map and event sparkline showing key phospho-events during insulin stimulation. Shows 24 events inferred from 17 clusters of 3,172 phosphosite profiles (Supplementary Fig. 3). **a**, Event map of key phosphorylation (up arrows) and dephosphorylation (down arrows) activation events. Each horizontal bar represents one cluster, and indicates when phosphorylation is increased. Each column (gray or white shading) indicates a group of events that occur at significantly different times to all other events, based on pairwise Mann–Whitney U tests (false discovery rate corrected). Events with the same horizontal positioning are assessed to occur simultaneously. Events connected with dotted gray lines occur at overlapping times (based on pairwise statistical tests); some of these events are nonetheless inferred to be non-simultaneous (based on graph traversal). **b**, Event sparkline summarizing the inferred temporal order of events shown in above event map. Events are labelled with cluster numbers, with multiple numbers indicating simultaneous events (e.g., the dephosphorylation events in clusters 1 and 4). Gray lines or arcs indicate overlapping but non-simultaneous events. Figures made using data from Humphrey *et al*.^*14*^ with Minardo-Model and edited in Adobe Illustrator.

**Fig. 3.**
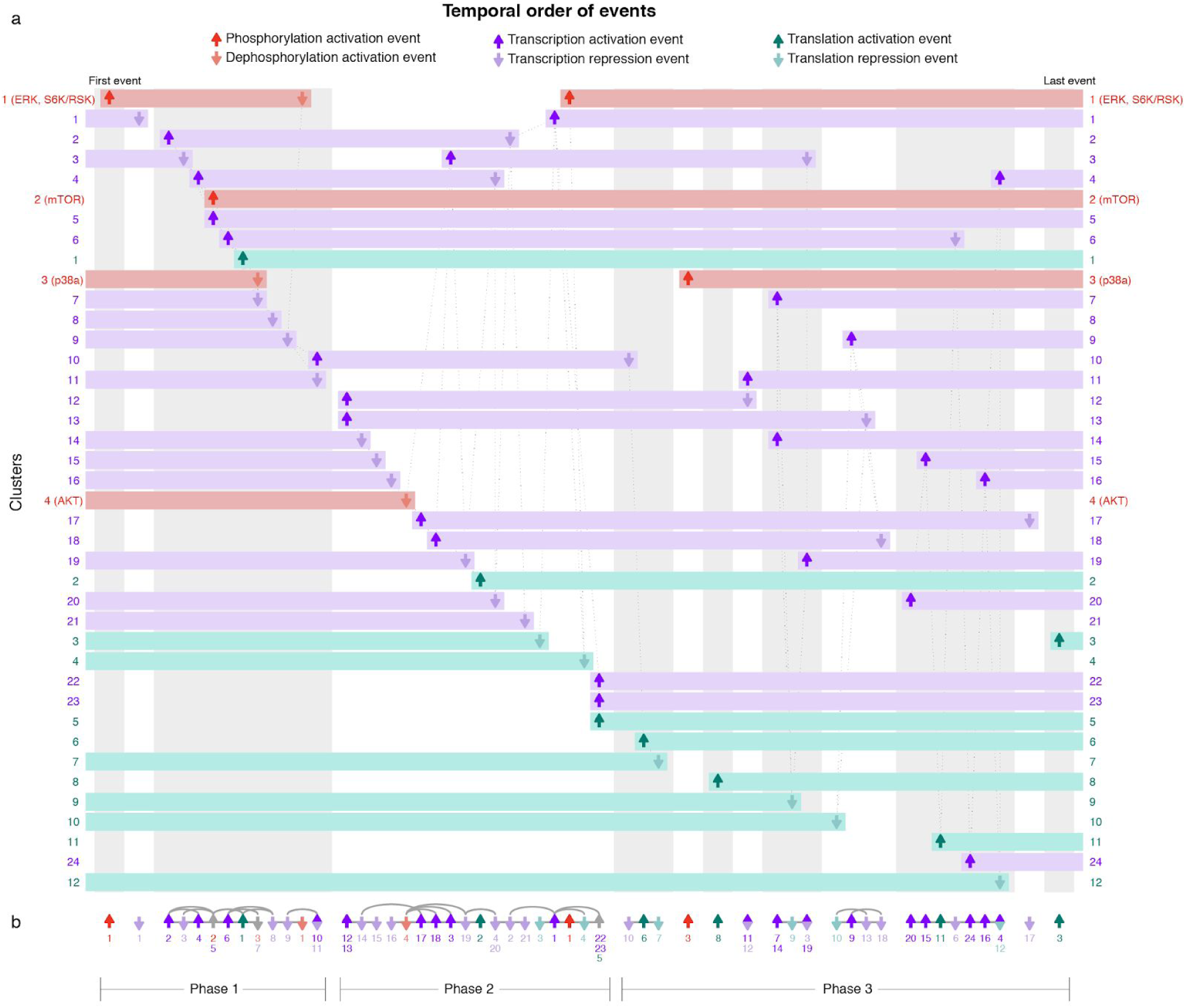
Event map and event sparkline showing multiomics events during epiblast differentiation. Shows 64 multiomics events inferred from 40 clusters: 4 clusters comprise 3,564 phosphosite profiles (red), 24 clusters comprise 6,225 mRNA profiles (purple), and 12 clusters comprise 2,735 protein profiles (green; see Supplementary Figs. 4-6). **a**, Event map of key phosphorylation, transcription, and translation events, with up and down arrows indicating activation and repression, respectively. Each horizontal bar represents one cluster, and indicates when it has increased abundance. Each column (gray or white shading) indicates a group of events that occur at significantly different times to all other events. Events with the same horizontal positioning are assessed to occur simultaneously. Dotted gray lines indicate overlapping but non-simultaneous events. **b**, Event sparkline summarizing the inferred temporal order of events shown in above event map. Events are labelled with cluster numbers. Gray lines or arcs indicate overlapping but non-simultaneous events. The large group of 22 overlapping events (‘Phase 2’) drive exit from the naive state; prior events (‘Phase ‘1) mostly relate to stem cell renewal; later events (‘Phase 3’) drive epiblast induction (see Supplementary note). Figures made using data from Yang *et al*.^*8*^ with Minardo-Model and edited in Adobe Illustrator.

The first test data set was from Humphrey *et al*. and measured the phosphoproteomic responses to insulin stimulation in mouse pre-adipocyte 3T3-L1 cells^14^. This experiment tracked changes over 48 hours at 9 time points (including basal), finding significant changes at 3,172 phosphosites, which were partitioned into 17 clusters^6^ using FCM^3^. Analysing these clusters using Minardo-Model, we found that the distribution of standardized abundances was always approximately normally distributed (Supplementary Fig. 2), thus justifying subsequent use of GLMs^9^. However, the distribution of event times was occasionally highly skewed (Supplementary Fig. 3), thus the non-parametric Mann-Whitney U test was used, resulting in an ordering for 24 inferred events (Fig. 2). The automatically derived ordering was found to be strongly correlated (Spearman’s ρ = 0.81^15^) with a previously published ordering derived manually via an extensive literature survey^1,2,7,16^ (see Supplementary note for details).

The second test data set was from Yang *et al*. and measured multiomic changes during differentiation of mouse E14Tg2a cells from the naive embryonic stem cell (ESC) state to primed epiblast-like cells (EpiLC)^8^. This experiment tracked changes in phosphorylation, RNA abundance, and protein abundance over 72 hours at 8 time points (common to each omics technique). Yang *et al*. reported 3,585 significantly changed phosphosites, partitioned into 4 clusters (Supplementary Fig. 4), and 6,225 mRNAs with significant changes in abundance, which we partitioned into 24 clusters using STEM^5^ (Supplementary Fig. 5). Yang *et al*. also reported 2,735 proteins with significant changes in abundance, partitioned into 12 clusters (Supplementary Fig. 6). We used Minardo-Model to analyse all clusters combined, resulting in 64 ordered, multiomic events concisely summarized into a single, integrated figure (Fig. 3). These events divided naturally into three phases: the phase 1 transcription and translation events were associated with stem cell renewal, while the phosphorylation events signalled the start of the next phase; phase 2 was associated with exiting the naive state; phase 3 was associated with entry into the primed epiblast state. The order and composition of these events agreed closely with the manual analysis from Yang *et al*. and with many other previous studies on epiblast differentiation (see Supplementary note for details).

As with many analyses of omics data, the above evaluation is highly selective, focusing on only the small subset of phosphosites, genes and proteins with well-understood roles. Typically, in omics experiments, much remains unreported and unknown regarding many of the observed changes. We believe that Minardo-Model may help improve this situation by clearly revealing what is and is not known, thus freeing researchers to focus on higher level questions, generate hypotheses, and uncover novel insights buried in complex data.

Minardo-Model has several advantages over previous work. To our knowledge, it is the first method to infer ordering of omics events at better temporal resolution than the experiment. It can take, as input, outputs generated from a wide range of existing clustering and analysis methods. The inferred events are then overlaid onto the widely-used matrix of profile plots (Fig. 1, Supplementary Figs 4-6), but where individual profile variation has been colored in a manner which avoids limitations of rainbow color maps^2,6^ used in some previous work^2,6^. The novel event map and event sparkline layouts are useful for revealing overall trends, while still providing sufficient detail (via cluster numbering and event ordering) to provide insight into the behavior of individual biomolecules. Event maps can be particularly helpful in revealing causal cascades, such as where expression of a transcription factor leads to its subsequent transcriptional activity. Event sparklines may be especially useful in temporal scRNAseq studies of complex differentiation processes, where the sparklines could be combined with currently used layouts, such as developmental trajectories, to provide greater detail – and potentially more insight – with relatively little increase in visual complexity. Additionally, sparklines may be useful to visually summarize differences obtained when data are analyzed using various clustering methods.

A potential limitation of our method is that, for larger and more noisy data sets, more events may be assessed as simultaneous: an example is shown in Fig. 2, where events in clusters 14 and 15 are simultaneous, so are placed in the same column. Also, more events may be assessed as non-simultaneous but overlapping: an example is shown again in Fig. 2, with a group of 6 overlapping events from clusters 2, 3, 10, 11, 12, and 13. In this case, based on the pattern of connections shown (which indicate event pairs with significant overlap in timing), Minardo-Model can infer the ordering 10/11 → 3 → 12 → 2/13 (Fig. 2b). However, for more complex data sets where as such situations can become exacerbated (e.g., Fig. 3), more powerful statistical approaches may be needed.

Another potential limitation is that the event ordering obtained depends on the criteria used. Currently, Minardo-Model uses somewhat arbitrary criteria to assign event timing, based on when the individual abundance profiles cross a 50% threshold – at which point approximately half of the changes induced by the hidden event will have occurred. Other criteria could be used – e.g., based on inflection points in the abundance profiles (Supplementary Fig. 3), or based on unbiased thresholding methods (e.g., Otsu^17^). Identifying optimal criteria for defining events could be a ripe topic for further research; practically, however, the major limitation will usually be the high variability and noise found in many current experimental data sets.

In summary, Minardo-Model provides a novel method for analysing and visualizing time series data sets from omics and multiomics experiments. Staring with input data partitioned using any preferred clustering method, Minardo-Model helps interpret the resulting clusters by automatically inferring an event ordering (often, at better temporal resolution than in the original experiment), and by providing concise, integrated visual summaries showing all underlying events in the data set (Fig. 2). This, in turn, can help in gaining insight into the processes underlying cellular responses and state changes^7^ (Fig. 3). Minardo-Model is freely available as an R package at https://bit.ly/MinardoModel.

## Methods

### Input data

Minardo-Model is designed to take as input a set of individual time profiles of abundance values, a set of clusters (calculated using any preferred method), and a matrix that assigns each individual profile a membership score for each cluster. In the cases used in this work, the membership scores were calculated via fuzzy c-means clustering (FCM^3^), however outputs from hard clustering methods can also be used. As a first step (Supplementary Fig. 1), each individual profile is standardized by subtracting the mean abundance value from the value at each time point, then dividing by the standard deviation.

### Abundance distributions

Next, Minardo-Model automatically generates a distribution of standardized abundances for each time point in each cluster (Supplementary Figs. 2 & 7). The plots are generated using the density function in the R ‘stats’ package, with default Gaussian kernel and bandwidths of 0.5 and 0.4, respectively. These distributions should be checked to ensure they are all approximately normally distributed; if not, before proceeding, the clustering strategy or input data needs to be changed until this step gives normal distributions, otherwise subsequent use of generalized linear models^9^ (GLM) would not be valid.

### Event windows

For each cluster, Minardo-Model then formulates a GLM^9^, using standardized abundances as a response variable determined by two predicting factors: time point and a unique identifier for each individual profile. For each pair of time points in the cluster, the GLM is then used to generate a Z-score, and a Tukey post-hoc test^10^ is used to calculate a p-value. Using default parameters, a pair of time points are considered to have a significant change in abundance only if |Z| > 15 and Tukey p < 0.001 (both parameters can be adjusted in Minardo-Model). A list of time point pairs with significant changes is then created and ranked by Z-score; the list is then used to assign event windows as follows:

1. Starting with the pair of time points that correspond to the largest Z-score, these time points are used to define a putative event window.
2. For each pair of consecutive time points within the putative event window, check whether each corresponding Z-score changes monotonically (i.e., always increasing or always decreasing).
3. If step 2 is true, then check that the putative event window does not overlap with any previously defined event window.
4. If steps 2 and 3 are both true, then a new event window is defined; if not, the putative window is rejected and no event window is defined using that Z-score.
5. Repeat steps 2-5 with putative event windows defined using progressively lower Z-scores, until all Z-scores have been considered.

Each event window defined in this way is assumed to arise due to one or more underlying events (e.g., the activation of a transcription factor, kinase, or phosphatase).

### Heat maps

For each cluster, Minardo-Model then generates 2D heat maps showing Z-scores between all pairs of time points (including non-consecutive time points) for each cluster. For brevity, the complete set of 2D heat maps are not shown here; instead, we show only a single row from each heat map, corresponding to the Z-scores between consecutive time points in each cluster (Fig 1).

### Event timing

Within each event window, an individual event time is then calculated for each individual profile, defined as the time at which the profile crosses halfway between the lowest and highest median abundance values within each event window, as estimated by linear interpolation (hereafter referred to as the 50% threshold). The distribution of individual event times are shown via density plots (Supplementary Figs. 3, 8-10), generated using the density function in the R ‘stats’ package, with default Gaussian kernel and bandwidths of 0.5 and 0.4, respectively. These density plots, in turn, are used to define either the mean or median event time (*t*_*0*.*5*_), for normal or skewed distributions, respectively. Optionally, the user can choose to set the threshold to a different value to suit their specific experimental scenarios.

### Profile plots

For all clusters, Minardo-Model then automatically generates a matrix of profile plots; each plot shows one cluster, including all individual profiles (thin transparent lines), cluster centroids (thick lines), events, and event windows (Fig. 1 & Supplementary Figs 4-6).

### Event ordering

The median or mean times calculated for each event implies an initial, or putative, ordering – however, in the following steps, this putative ordering is refined using statistical and graphical methods, resulting in a final ordering supported by all available evidence.

1. For each event window, a t_0.5_ value is calculated for each individual profile, resulting in a distribution of individual event times.
2. The t_0.5_ values are then plotted as a series of density plots (Supplementary Figs. 3, 8-10) that should be checked to determine whether they are all approximately normally distributed.
3. If the events are normally distributed, Minardo-Model should then be set to use Student’s t-test^11^ for comparing distribution pairs; otherwise, the comparisons should be done using Mann-Whitney U tests^12^.
4. From the above comparisons, two matrices are computed: one matrix (M) stores the test statistics (i.e., Student’s t scores or Mann-Whitney’s U scores) and the other matrix (P) stores corresponding p-values, adjusted via a false discovery rate (FDR) correction^18^. Each of these matrices will be of size n×n, where n is the number of event windows.
5. From M and P, a new Boolean matrix is computed, such that B_i,j_ = 1 if i ≠ j, P_i,j_ < 0.05, and M_i,j_ < 0; otherwise B_ij_ = 0, where i and j represent events. Here, B is similar to an adjacency matrix for a directed graph, where the nodes now represent events, and a directed edge between two nodes indicates that the source event occurs significantly earlier than the target event. Pairs of nodes with no connecting edge indicate events with no statistically significant difference in timing – hereafter referred to as overlapping events. Thus, we have represented events and relationships between events as a directed graph.
6. B is then transitively reduced^19^, resulting in a graph containing a minimal number of edges (B′).
7. B′ is then used to calculate a matrix (O) that maps events to a unique ordering. O is determined by traversing B′ via a breadth-first search algorithm^20^, such that at any depth (d) in the search tree, all events are overlapping, whereas for any two adjacent depths (d and d+1), at least one event at depth d occurs significantly earlier than at least one event at d+1. Note that the root node of the breadth first search graph (B′) corresponds to the earliest events.

As a result of the above steps, all events with the same rank in the ordering will overlap with each other (i.e., have no significant differences in timing based on pairwise statistical comparisons); these are hereafter referred to as simultaneous events, and they are visualized in a manner that indicates they occur simultaneously (e.g., in Fig. 2, the first event in cluster 14 and the last event in cluster 15).

Note that some overlapping events are not simultaneous in the ordering derived by graph traversal; as described below, such cases are highlighted using undirected edges in the event map and event sparkline (e.g., in Fig. 2, the first event in cluster 5 and the last events in clusters 1, 4, and 6). For brevity, these are the only edges drawn between events.

We also note that the final ordering (O) can differ from the initial, putative order based on the mean event times. For this reason, in the subsequent plots, the layout of events is designed to show event order, not the mean timing of events.

### Event maps

Next, Minardo-Model automatically plots an event map using the following steps:

1. The ordering (O) of the first event in each cluster is used to sort all clusters vertically, with the first event (and corresponding cluster) placed top-left.
2. Next, all ordered events are partitioned into a series of event groups, defined as a series of consecutively ordered events that contain some overlaps, based on pairwise statistical comparison. Each event group is visually indicated as a separate column, using alternating background shading, and by adding a slight horizontal spacing.
3. Within each column, all overlapping events are connected with undirected edges, drawn as dashed gray lines.
4. Events involving an increase in abundance (e.g., gene expression) are indicated using bright red, upward arrows, while decreasing events (e.g., gene repression) are indicated with dull red, downward arrows.
5. Finally, each cluster is drawn as a series of rectangles, beginning and ending with events, thereby showing times when profiles in the cluster are upregulated.

### Event sparklines

Finally, Minardo-Model plots an event sparkline; this is a similar albeit simpler process than plotting event maps. Here, the ordering of events (O) is directly indicated horizontally. Each event group is visually indicated by adding a slight horizontal spacing, and by connecting all non-simultaneous but overlapping events with undirected edges, drawn as gray lines. These lines are drawn as arcs for overlapping events that are not adjacent in the ordering (O) derived via graph traversal (Figs 2b & 3b).

### Phosphoproteomics data set

We tested our method by applying it to a time-series data set published by Humphrey *et al*.^*14*^, which measured global cellular phosphoproteomic changes in response to insulin stimulation at 9 time points (including basal) in 3T3-L1 cells. In total, 37,248 phosphorylation sites were detected, of which 3,172 were assessed to show significant change in response to insulin. To use this data set, we first needed to repeat the following 4 analysis steps done by Humphrey et al.^14^, since the published data set lacks some of the intermediate, processed data they reported. First, we identified the significantly changed profiles by performing empirical Bayes moderated t-tests (using the limma R package^21^) with significance defined when the false-discovery rate (FDR) ≤ 0.05 and a fold change of 2. These significantly changed profiles were then standardized (see above) and partitioned into 17 clusters using the FCM^3^ algorithm in the Mfuzz R package^22^ (Supplementary Table 1). The partitioning into 17 clusters was reported to be optimal by Yang *et al*.^*6*^ for this data set.

### Multiomics data set

We also tested Minardo-Model by applying it to a multiomics data set published by Yang *et al*.^*8*^, which measured responses in mouse E14Tg2a cells as they differentiated from the naive embryonic stem cell (ESC) state to primed epiblast-like cells (EpiLC)^8^. Yang *et al*. measured phosphoproteomics changes at 12 time points (including basal) and provided 3,585 phosphosite profiles partitioned into 4 clusters using CLUE^6^. We removed 21 profiles, since their corresponding protein name was not provided, and the remaining 3,564 profiles were analysed with Minardo-Model (Supplementary Fig. 4).

Yang *et al*. also measured transcriptomic changes at 8 time points (including basal), where a total of 20,936 transcripts were quantified at all time points. These data were not clustered by Yang *et al*. Hence, first, we identified the significantly changed profiles, in a manner similar to the phosphoproteomics Humphrey *et al*. data set (see above), by performing empirical Bayes moderated t-tests (using the ‘limma’ R package^21^) with significance defined when the false-discovery rate (FDR) ≤ 0.05 and a fold change of 2. This resulted in a total of 7,945 profiles. These significantly changed profiles were then standardized and clustered using the Short Time-series Expression Miner (STEM) tool^5^. To STEM, we provided as input the 7,945 normalized gene expression profiles, without the basal value. We then chose the option ‘No normalization/add 0’ and did not provide any annotation sources. Finally, we set the ‘Maximum number of model profiles’ to 139. We found this number to be optimal, since specifying a larger number resulted in clusters being split across them. We then ran STEM using default values for all remaining settings, which gave 24 clusters containing a significant number of genes (Supplementary Table 3). A total of 6,225 genes were included in these 24 clusters (Supplementary Fig. 5) and utilized for further analysis with Minardo-Model.

Finally, Yang *et al*. also measured proteomic changes at 9 time points (including basal) and provided 2,735 profiles clustered into 12 clusters using FCM^3^. Six of these clusters contained up-regulated proteins and six contained down-regulated proteins (Supplementary Fig. 6). We utilized these clusters directly for analysis with Minardo-Model.

Using Minardo-Model, the phosphoproteomics, transcriptomics, and proteomics data were initially analyzed independently, and then combined during the final event ordering step. We found that the distributions of standardized abundances were always approximately normally distributed (see Supplementary Fig. 7), thus justifying the subsequent use of GLMs^9^. The distribution of event times was occasionally highly skewed (Supplementary Figs. 8-10), so the non-parametric Mann-Whitney U test was then used to do the final pairwise comparisons of event times, combining events from all three subsets of the multiomics data (Fig 3).

## Acknowledgements

JRK is a recipient of the Australian Diabetes Society Skip-Martin Early Career Fellowship. We gratefully acknowledge Martin Krzywinski for useful discussions. We are also grateful to the anonymous reviewers of this manuscript, whose detailed and insightful feedback helped to make significant improvements.

## Author contributions

SK and SIO conceived the idea. SK performed all the analysis, wrote code and determined the algorithm for ordering events. SK and SIO wrote the manuscript and created the figures. SK, TJP and SIO identified strategy for identifying and comparing events and defining event windows. TJP identified strategy for analyzing change within clusters using GLMs. LDWL and SK evaluated the multiomic data set ordering. PY provided data sets for analysis as well as code for identifying differentially altered profiles and standardizing them. JRK provided useful discussions regarding all aspects of the work. JV tested the R package. SIO guided research and performed all supervisory tasks.

## References

1. Kaur, S., Baldi, B., Vuong, J. & O’Donoghue, S. I. Visualization and analysis of epiproteome dynamics. J. Mol. Biol. 431, 1519–1539 (2019).

2. O’Donoghue, S. I. et al. Visualization of Biomedical Data. Visualization of Biomedical Data (2018).

3. Bezdek, J. C., Ehrlich, R. & Full, W. FCM: The fuzzy c-means clustering algorithm. Computers & Geosciences 10, 191–203 (1984).

4. Bar-Joseph, Z., Gitter, A. & Simon, I. Studying and modelling dynamic biological processes using time-series gene expression data. Nat. Rev. Genet. 13, 552–564 (2012).

5. Ernst, J. & Bar-Joseph, Z. STEM: a tool for the analysis of short time series gene expression data. BMC Bioinformatics 7, 191 (2006).

6. Yang, P. et al. Knowledge-Based Analysis for Detecting Key Signaling Events from Time-Series Phosphoproteomics Data. PLoS Comput. Biol. 11, e1004403 (2015).

7. Ma, D. K. G., Stolte, C., Krycer, J. R., James, D. E. & O’Donoghue, S. I. SnapShot: Insulin/IGF1 Signaling. Cell 161, 948–948.e1 (2015).

8. Yang, P. et al. Multi-omic Profiling Reveals Dynamics of the Phased Progression of Pluripotency. Cell Syst. 8, 427–445.e10 (2019).

9. McCullaugh, P. M. & Nelder, J. A. Generalized linear models 2nd edition. (1989).

10. Tukey, J. W. Comparing individual means in the analysis of variance. Biometrics 5, 99–114 (1949).

11. Student. The Probable Error of a Mean. Biometrika 6, 1 (1908).

12. Mann, H. B. & Whitney, D. R. On a test of whether one of two random variables is stochastically larger than the other. Ann. Math. Statist. 18, 50–60 (1947).

13. Tufte, E. R. Beautiful evidence. 1, (Graphics Press LLC, Cheshire, 2006).

14. Humphrey, S. J. et al. Dynamic adipocyte phosphoproteome reveals that Akt directly regulates mTORC2. Cell Metab. 17, 1009–1020 (2013).

15. Spearman, C. The proof and measurement of association between two things. Int. J. Epidemiol. 39, 1137–1150 (2010).

16. Kaur, S., Baldi, B., Vuong, J. & O’Donoghue, S. I. A benchmark dataset for analyzing and visualizing the dynamic epiproteome. Data Brief 25, 104000 (2019).

17. Otsu, N. A Threshold Selection Method from Gray-Level Histograms. IEEE Trans. Syst. Man Cybern. 9, 62–66 (1979).

18. Benjamini, Y. & Hochberg, Y. Controlling the false discovery rate: A practical and powerful approach to multiple testing. Journal of the Royal Statistical Society: Series B (Methodological) 57, 289–300 (1995).

19. Aho, A. V., Garey, M. R. & Ullman, J. D. The Transitive Reduction of a Directed Graph. SIAM J. Comput. 1, 131–137 (1972).

20. Skiena, S. S. The algorithm design manual: Text. 1, (Springer Science & Business Media, 1998).

21. Smyth, G. K., Thorne, N. & Wettenhall, J. Limma: linear models for microarray data user’s guide. Software manual available from http://www.bioconductor.org (2003).

22. Kumar, L. & E Futschik, M. Mfuzz: a software package for soft clustering of microarray data. Bioinformation 2, 5–7 (2007).

