## Supplementary_information for "Temporal ordering of omics and multiomic events inferred from time series data"

#### Additional files

Supplementary Table 1 - Phosphoproteomics data set comprising 3,172 phosphosite profiles partitioned into 17 clusters along with event timings derived by Minardo-Model.
Supplementary Table 2 - Comparison of phospho-event ordering derived by Minardo-Model with manual ordering from Cell SnapShot.
Supplementary Table 3 - Multiomics data set comprising 3,585 phosphosite profiles partitioned into 4 clusters, 6,225 mRNA profiles partitioned into 24 clusters, and 2,735 protein profiles partitioned into 12 clusters along with along with event timings derived by Minardo-Model.

Supplementary code R package: Minardo-Model hosted at: https://bit.ly/MinardoModel.

#### Supplementary figures

**Flowchart depicting the typical workflow using Minardo-Model**

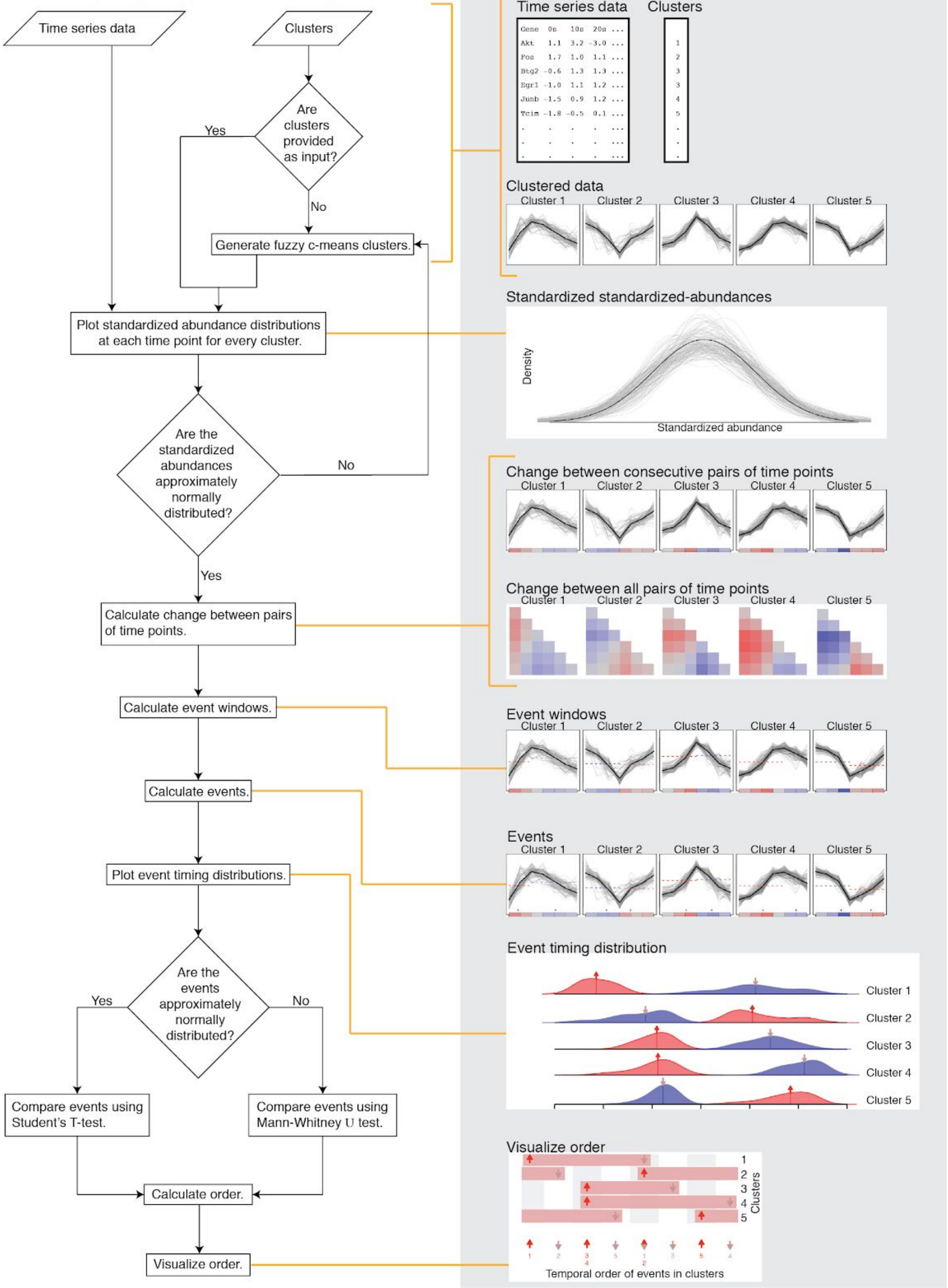

**Supplementary Fig. 1 | Minardo-Model workflow with example outputs.** Minardo-Model takes time-series data as input, as well as optional cluster assignments for each profile. If cluster assignments are missing, they are generated using FCM[1]. For each cluster, Minardo-Model then calculates the following: (a) a distribution plot showing abundances at each time point, allowing the user to assess whether these data are normally distributed. If not, the clusters must be redone, or the data modified, before proceeding. Next, Minardo-Model calculates a matrix showing abundance changes between all pairs of time points; these data are then used to calculate event windows, and events. The distribution of events is plotted, and if a normal distribution is followed, then Student's t-test can be utilized, otherwise the slower Mann-Whitney U test can be utilized for subsequent event distribution comparison and event ordering. Finally, the event ordering is visualized using event maps and sparklines.

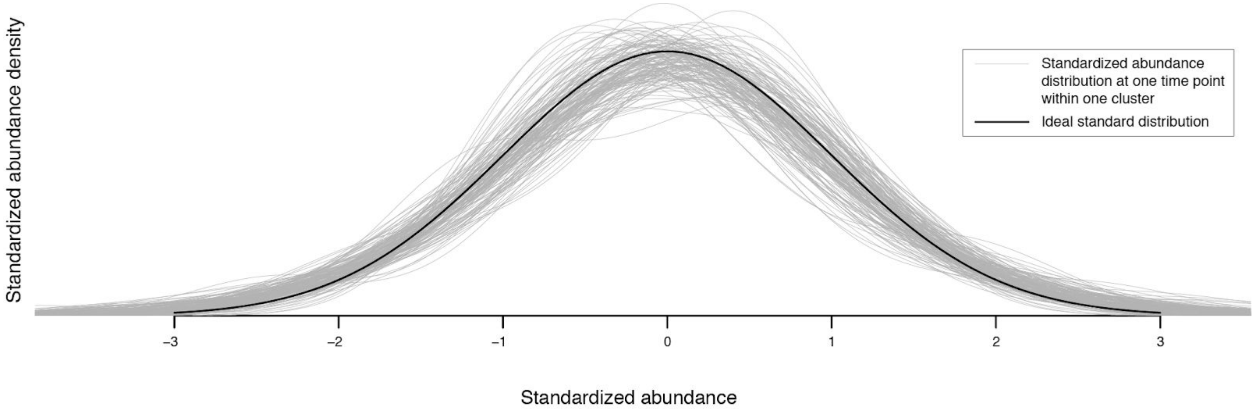


**Supplementary Fig. 2 | Standardized abundance distributions from the phosphoproteomics data set.** Shows a superposition of 153 standardized abundance distributions (thin gray lines), one for each of the 9 time points within each of the 17 clusters of the phosphoproteomics data set. Each distribution was calculated by scaling the abundance values at one time point from all the individual profiles within a cluster (setting the mean value to 0 and standard deviation to 1), then generating a Gaussian kernel density plot. For comparison, the ideal normal distribution for these parameters is shown (thick black line). The figure shows that, for this data set, the abundance distributions are all approximately normally distributed, thus justifying subsequent use of GLMs. Figures made using data from Humphrey et al.[2] with Minardo-Model.

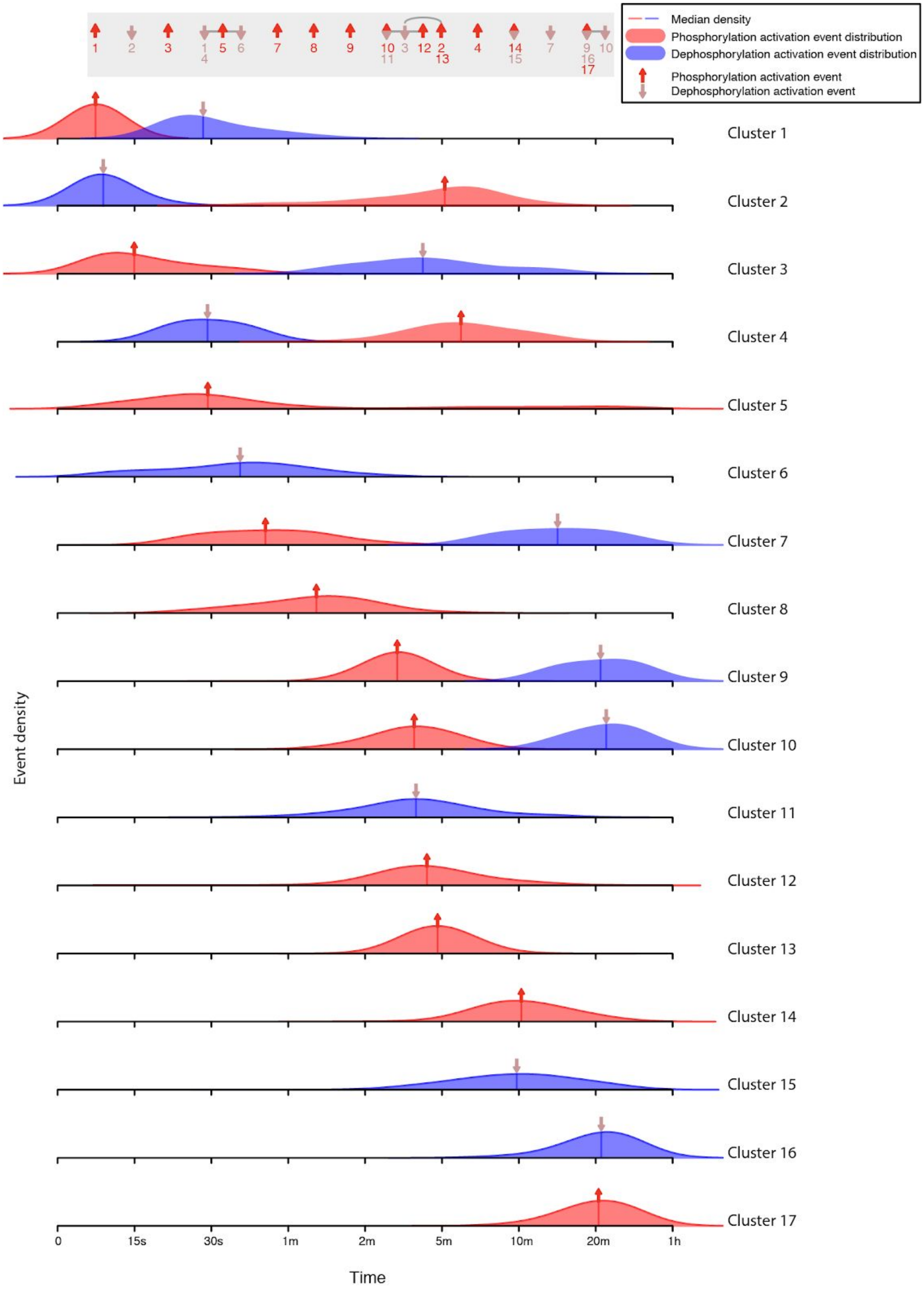

Median density
Phosphorylation activation event distribution
Dephosphorylation activation event distribution
Phosphorylation activation event
Dephosphorylation activation event
Cluster 1
Cluster 2
Cluster 3
Cluster 4
Cluster 5
Cluster 6
Cluster 7
Cluster 8
Cluster 9
Cluster 10
Cluster 11
Cluster 12
Cluster 13
Cluster 14
Cluster 15
Cluster 16
Cluster 17
Event density
0
15s
30s
1m
2m
5m
10m
20m
1h
Time

**Supplementary Fig. 3 | Distribution of event times in the phosphoproteomics data set.** Shows the distribution of event times calculated from each individual profile in each cluster. Each kernel density plot has been scaled to have the same total area under the curve, and the median event time is indicated via an arrow and a vertical bar. Some distributions are skewed or bimodal, thus the non-parametric Mann-Whitney $U$ test was used to statically assess the ordering of the median events. For reference, the final derived event ordering is shown, at the top of the graph, as an event sparkline (from Fig. 2). Figures made using data from Humphrey et al.[2] with Minardo-Model and edited with Adobe Illustrator.

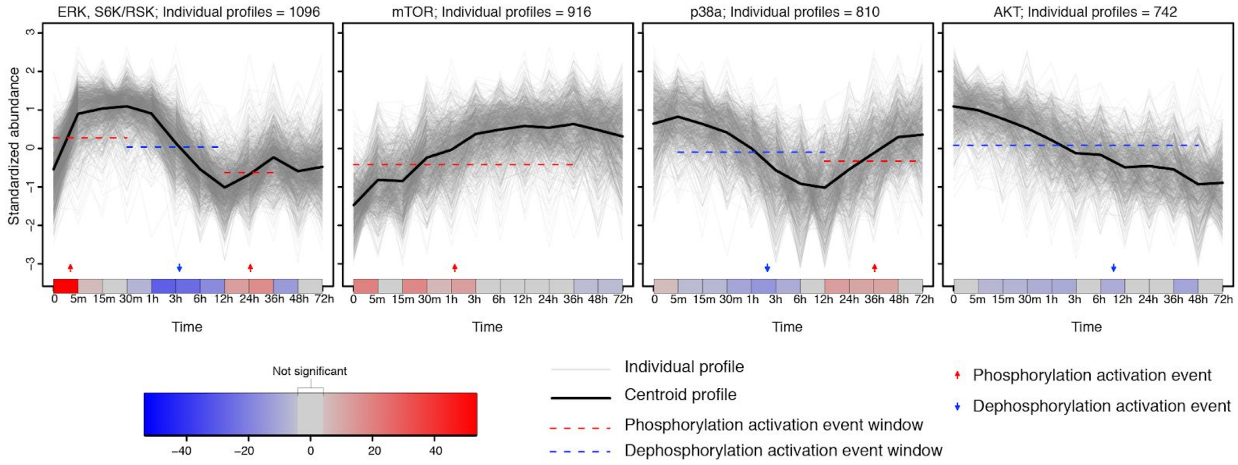


**Supplementary Fig. 4 | Profile plots of phosphorylation changes in a multiomics study of stem cell differentiation.** Individual profiles for 3,564 phospho-sites grouped into 4 clusters via CLUE[3], taken from Yang *et al.*[4]. Below each cluster, a single-row heat map indicates significant changes in mean phosphopeptide abundance between consecutive time points (red and blue showing phosphorylation and dephosphorylation, respectively), calculated via generalized linear models derived from individual profiles, and using Z-scores from a post-hoc Tukey test. The Z-scores and p-values are used to compute event windows, within which events are defined. Thus, the behavior of each cluster is summarized as a series of discrete phosphorylation and dephosphorylation activation events (red and blue arrows, respectively), based on the median time at which all individual profiles in a cluster cross half-maximal abundance within each event window (identified by the red or blue dashed line, respectively, shown here at the half maximal abundance of the centroid) - see Supplementary Fig. 8 for details. The cluster centroids are shown only to provide a graphical indication of the trend within each cluster. Figures made using data from Yang et al.[4] with Minardo-Model and edited with Adobe Illustrator.

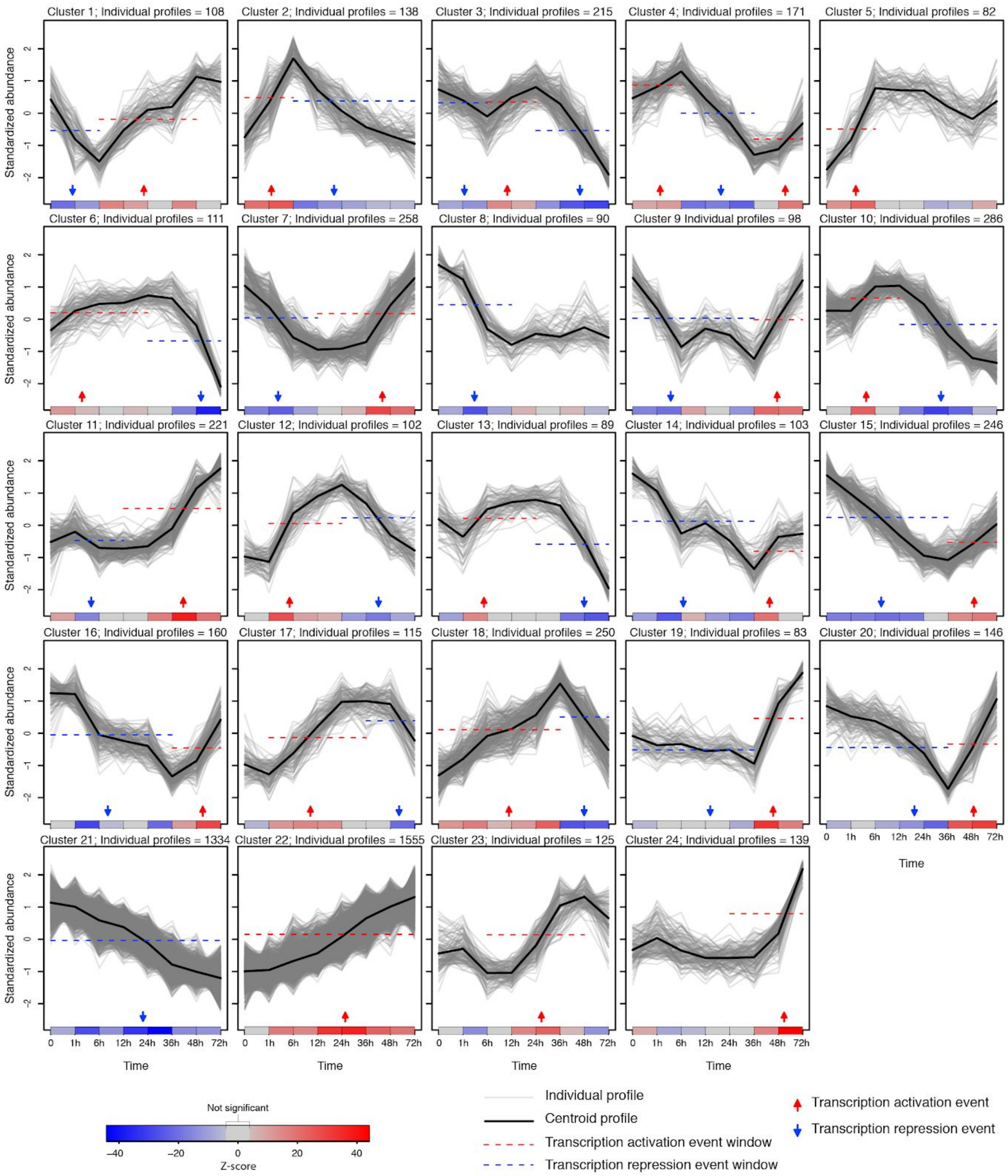


**Supplementary Fig. 5 | Profile plots of transcriptomic changes in a multiomics study of stem cell differentiation.** Individual profiles for 6,225 genes grouped into 24 clusters using STEM[5]. Below each cluster, a single-row heat map indicates significant changes in mean transcriptomic abundance between consecutive time points (red and blue showing up and down regulation, respectively), calculated via generalized linear models derived from individual profiles, and using Z-scores from a post-hoc Tukey test. The Z-scores and p-values are used to compute event windows, within which events are defined. Thus, the behavior of each cluster is summarized as a series of discrete transcription activation and repression events (red and blue arrows, respectively), based on the median time at which all individual profiles in a cluster cross half-maximal abundance within each event window (identified by the red or blue dashed line, respectively, shown here at the half maximal abundance of the centroid) - see Supplementary Fig. 9 for details. The ordering of these events was determined statistically, then used to sort and number the clusters. The cluster centroids are shown only to provide a graphical indication of the trend within each cluster. Figures made using data from Yang et al.[4] with Minardo-Model and edited with Adobe Illustrator.

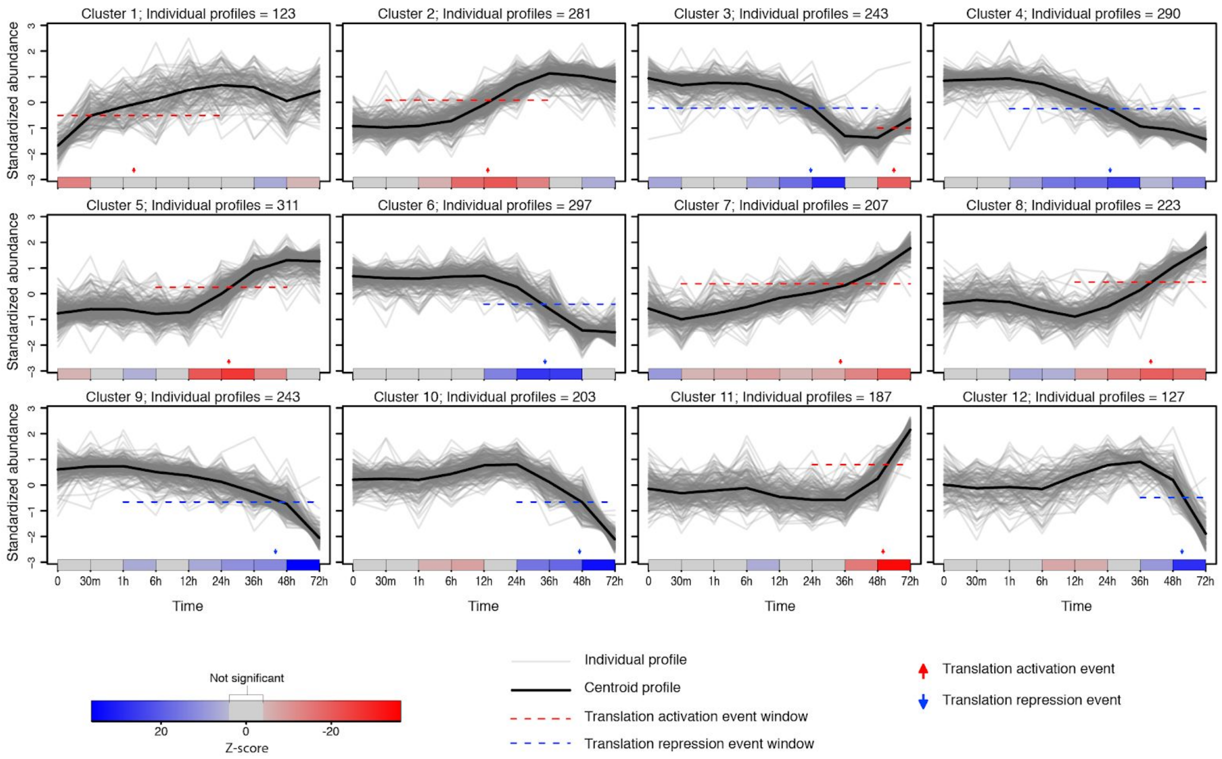


**Supplementary Fig. 6 | Profile plots of proteomic changes in a multiomics study of stem cell differentiation.** Individual profiles for 2,735 proteins grouped into 12 clusters via FCM[1], taken from Yang *et al.*[4]. Below each cluster, a single-row heat map indicates significant changes in mean protein abundance between consecutive time points (red and blue showing up and down regulation, respectively), calculated via generalized linear models derived from individual profiles, and using Z-scores from a post-hoc Tukey test. The Z-scores and p-values are used to compute event windows, within which events are defined. Thus, the behavior of each cluster is summarized as a series of discrete translation activation and repression events (red and blue arrows), based on the median time at which all individual profiles in a cluster cross half-maximal abundance within each event window (identified by the red or blue dashed line, respectively, shown here at the half maximal abundance of the centroid) - see Supplementary Fig. 10 for details.The ordering of these events was determined statistically, then used to sort and number the clusters. The cluster centroids are shown only to provide a graphical indication of the trend within each cluster. Figures made using data from Yang et al.[4] with Minardo-Model and edited with Adobe Illustrator.

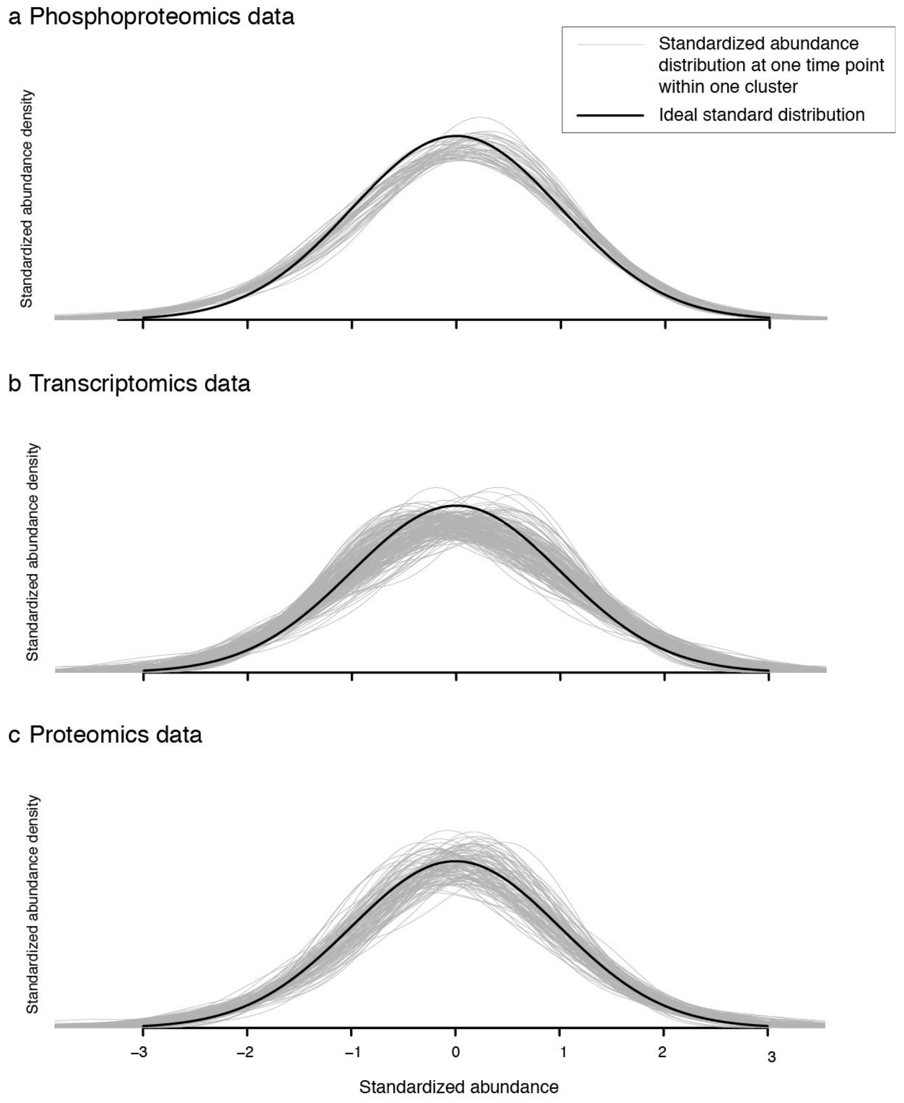


**Supplementary Fig. 7 | Standardized abundance distributions for the multiomics data set.** Each graph shows a superposition of standardized abundance distributions (thin gray lines), one for each of the time points within each of the clusters in the multiomics data set. Each distribution was calculated by scaling the abundance values at one time point from all the individual profiles within a cluster (setting the mean value to 0 and standard deviation to 1), then generating a Gaussian kernel density plot. For comparison, the ideal normal distribution for these parameters is shown (thick black line). The figure shows that, for this data set,  the abundance distributions at each time point are all approximately normally distributed, thus justifying subsequent use of GLMs. a, Shows 48 standardized abundance distributions, one for each of the 12 time points within each of the 4 clusters in the phosphoproteomics data. b, Shows 192 standardized abundance distributions, one for each of the 8 time points within each of the 24 clusters of the transcriptomics data. c, Shows 108 standardized abundance distributions, one for each of the 9 time points within each of the 12 clusters of the proteomics data. Figures made using data from Yang et al.[4] with Minardo-Model.

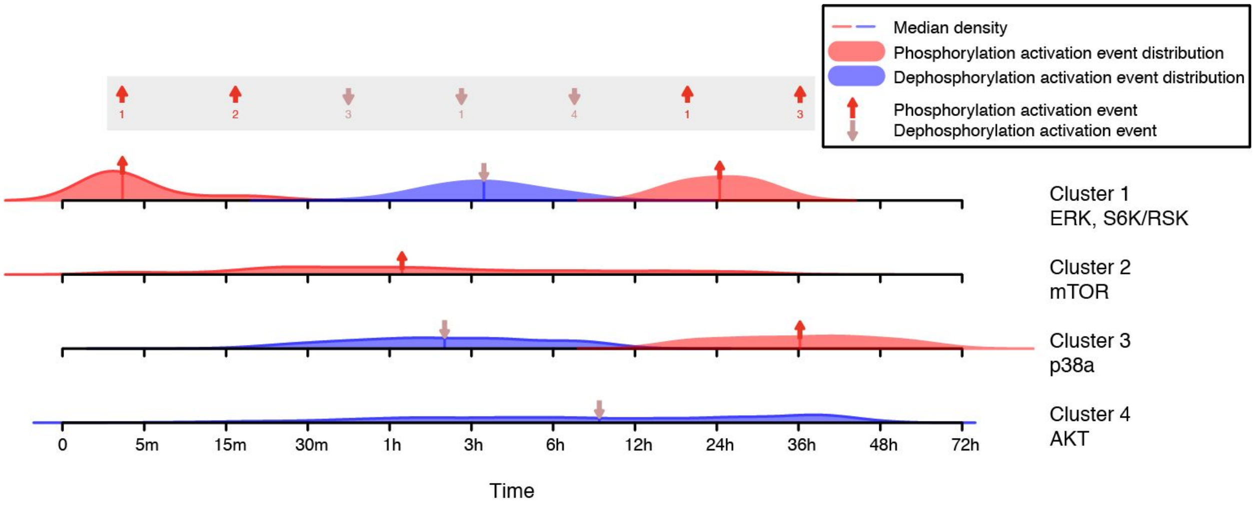


**Supplementary Fig. 8 | Distribution of phosphorylation event times in the multiomics data set.** Shows the distribution of event times calculated from each individual profile in each of the 4 phosphoproteomic clusters reported by Yang *et al.*[4] Each kernel density plot has been scaled to have the same total area under the curve, and the median event time is indicated via an arrow and a vertical bar. Some distributions are skewed or multimodal, thus the non-parametric Mann-Whitney U test was used to statistically assess the ordering of the median events. For reference, the final derived event ordering is shown, at the top of the graph, as an event sparkline (from Fig. 3). The very broad distributions seen in clusters 2-4 suggest that alternative clustering methods might be needed to describe the range of events occurring in this experiment. Nonetheless, the results reported by Yang *et al.* suggests that these four clusters may provide the best interpretation consistent with the data, due to the high levels of noise and variability in the experiment (see Supplementary Fig. 4). Figures made using data from Yang *et al.*[4] with Minardo-Model and edited with Adobe Illustrator.

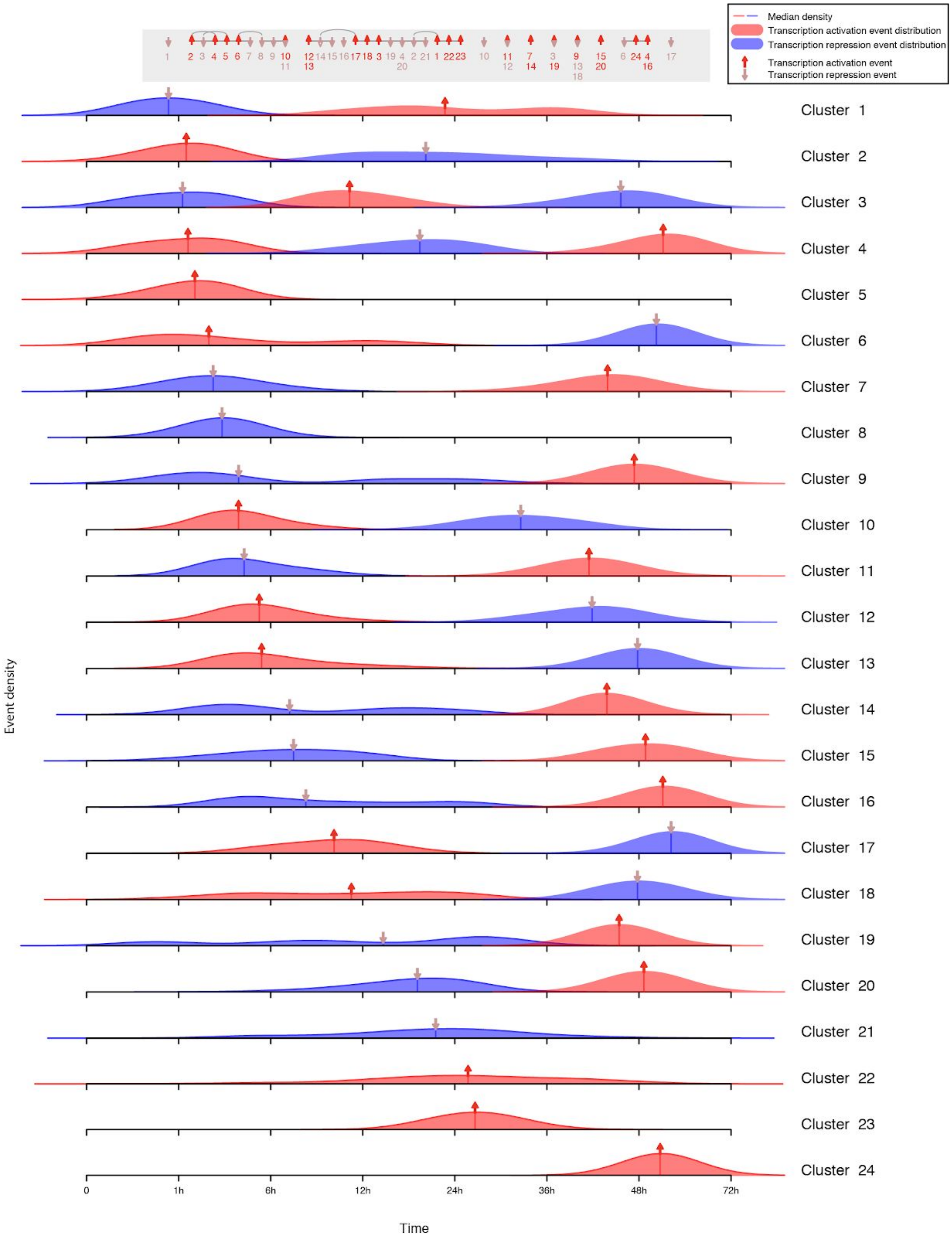


**Supplementary Fig. 9 | Distribution of transcription event times in the multiomics data set.** Shows the distribution of event times calculated from each individual profile in each of the 24 transcriptomic clusters calculated using STEM[5]. Each kernel density plot has been scaled to have the same total area under the curve, and the median event time is indicated via an arrow and a vertical bar. Some distributions are skewed or bimodal, thus the non-parametric Mann-Whitney $U$ test was used to statistically assess the ordering of the median events. For reference, the final derived event ordering is shown, at the top of the graph, as an event sparkline (from Fig. 3). The very broad distributions seen in clusters 2, 9, 14, 16, 18 and 19 suggest that alternative clustering methods might be needed to describe the range of events occurring in this experiment. However, the STEM analysis suggests that these may be the optimal clusters, given the high levels of noise and variability (see Supplementary Fig. 5). Figures made using data from Yang et al.[4] with Minardo-Model and edited with Adobe Illustrator.

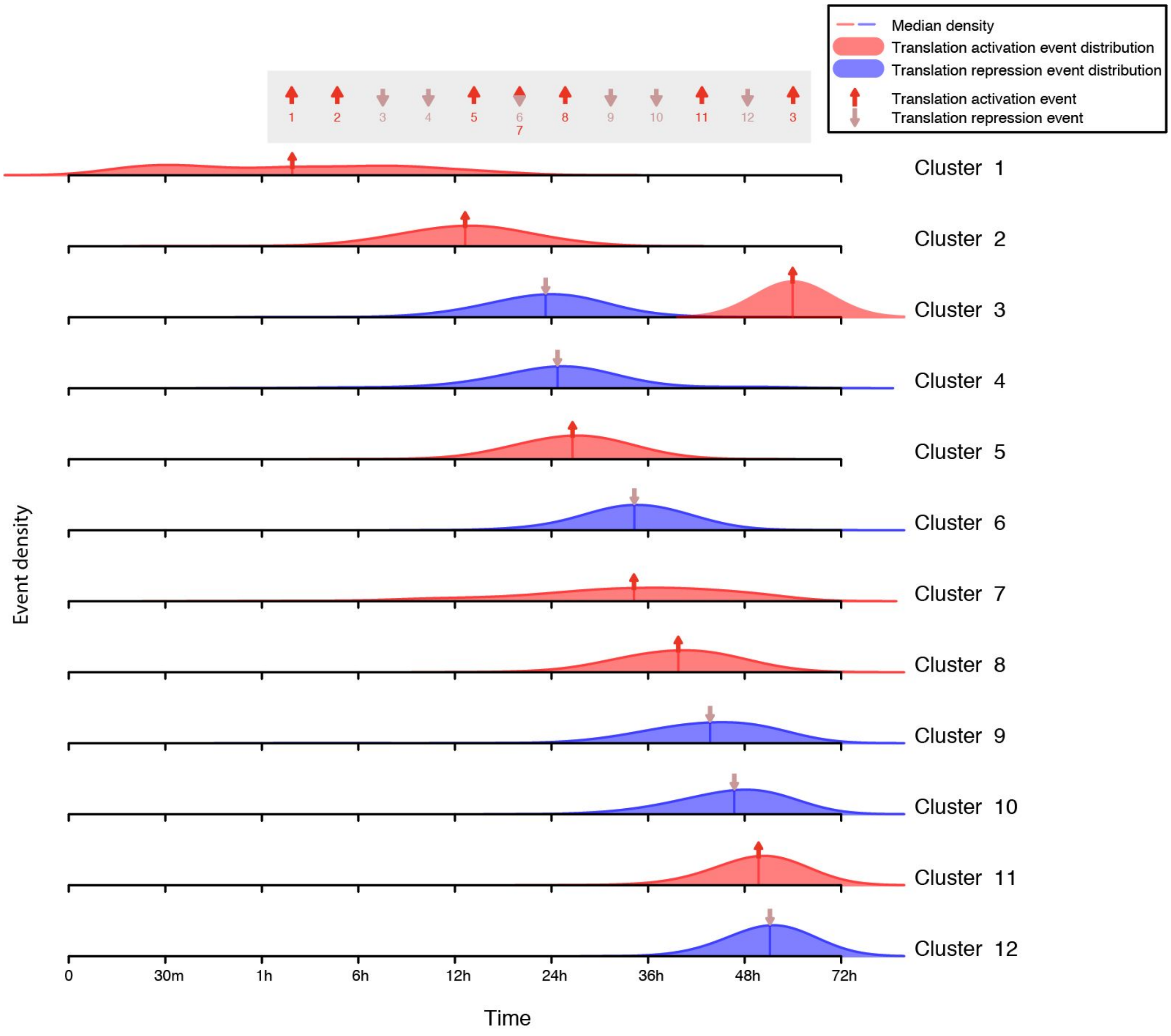


**Supplementary Fig. 10 | Distribution of translation event times in the multiomics data set.** Shows the distribution of event times calculated from each individual profile in each of the 12 proteomic clusters reported by Yang et al.[4] Each kernel density plot has been scaled to have the same total area under the curve, and the median event time is indicated via an arrow and a vertical bar. Some distributions are skewed or bimodal, thus the non-parametric Mann-Whitney U test was used to statistically assess the ordering of the median events. For reference, the final derived event ordering is shown, at the top of the graph, as an event sparkline (from Fig. 3). Figures made using data from Yang et al.[4] with Minardo-Model and edited with Adobe Illustrator.

### Supplementary notes

#### Evaluation of phosphoproteomics event ordering

We compared the order of events generated by Minardo-Model to the ordering for 103 key phospho-events in insulin/IGF1 signaling, derived from an extensive literature survey and previously published as a Cell SnapShot[6] and a Data in Brief[7].

We found a total of 48 phospho-events that were present in both the SnapShot and in 12 of the 17 clusters in the Humphrey *et al.* data set[2]. Of these, we removed two phospho-events (involving RPS6 S236 and S240) that were inconsistently annotated (dephosphorylated only in the SnapShot but phosphorylated only in the Humphrey *et al.* data set). We then compared the remaining 46 phospho-events using the 'stats' R package, finding a Spearman's correlation[8] of 0.81, confirming that the automatically-derived order identified by Minardo-Model is significantly correlated ($S = 3067$, $p < 10^{-11}$) with the manually-derived order of phospho-events reported in the SnapShot[6]. Further details of this comparison are included in Supplementary Table 2.

#### Evaluation of multiomics event ordering

Using data from Yang *et al.*, which tracked differentiation of mouse E14Tg2a from the naive embryonic stem cell (ESC) state to primed epiblast-like cells (EpiLC)[4], Minardo-Model inferred a temporal ordering for 60 multiomic events. Of these, 7 events were from four phosphoproteomic clusters (Supplementary Fig. 4), 13 events from 12 proteomic clusters (Supplementary Fig. 6), and 40 events from 24 RNA clusters (Supplementary Fig. 5). Following the analysis of Yang *et al.* these clusters were generated using CLUE[3] for the phosphoproteomics data and FCM[1] for the proteomics data. However, as Yang *et al.* did not provide clustering for the RNA data, we considered using several clustering methods (FCM[1], DREM[9,10]), ultimately deciding that STEM[5] was best suited to this experimental scenario. The resulting ordering of multiomic events is presented in Fig. 3, where we observed that, midway through the experiment, there was a natural grouping of 22 overlapping events (called 'phase 2'), beginning with transcriptional activation in cluster 12 and ending with translational repression in cluster 5. We used this grouping to partition all events into three phases: phase 1 comprised events prior to cluster 12 transcriptional activation, and phase 3 comprised events after cluster 5 translational repression.

We found that some of the events in phase 1 were associated with initiating the signalling cascade that heralds exit from ESC self-renewal processes[3,11]. These events include activation of kinases such as ERK, S6K, and mTor (phosphorylation clusters 1 and 2). However, the majority of events observed in this phase were found to be associated with maintenance of ESC self-renewal. This was evident in the repression of genes involved in Ras[12] and Notch signalling pathways[13] (RNA clusters 3 and 1, respectively), identified through KEGG pathway enrichment analysis (using the DAVID functional annotation tool[14] version 6.8). Additionally, we also observed repression of pathways associated with lineage determination, such as the 'hematopoietic cell lineage' (RNA cluster 8) and 'neurotrophin signalling' (RNA clusters 7 and 9).

By contrast, we found that most events in phase 2 were associated with exiting from ESC self-renewal. This was evident through the repression of key pluripotency transcription factors associated with the naive state[15,16], including Klf4, Klf2, Nr5a2, Prdm14, and Sox2 (in RNA cluster 21) as well as Nanog and Stat3 (in RNA cluster 21 and protein cluster 4). Additionally, phase 2 had events associated with epiblast induction, especially the activation of genes such as Sox3, Sox4, and Dnmt3b[17] (RNA cluster 22), as well as Otx2[18] (RNA cluster 22 and protein cluster 5).

Finally, we found that the events in phase 3 were associated with transition into the primed epiblast state; also, many of the transcriptional events observed in phase 2 associated with epiblast induction, manifested as translational events in phase 3. This included repression of Nr5a2, Nr0b1, Sox2, and Klf2 (protein cluster 7) and activation of Sox3 and Dnmt3b (protein cluster 6). In summary, our analysis showed that phosphorylation initiates the signalling cascade in phase 1, which was then reflected in transcriptional changes in phase 2, and in translational changes in phase 3, resulting in ESC to EpiLC transition.

We then compared the automated derived ordering inferred by Minardo-Model with the manual analysis of events for the proteome data reported by Yang *et al.*[4]. Yang *et al.* identified six activation and six repression events in the 12 clusters derived from the proteomics data subset. Minardo-Model found the same 12 events, plus one additional translation activation event in protein cluster 3, at approximately 60hr (Supplementary Table 3). Yang *et al.* also found that the proteomic changes occurred in three distinct waves.

Yang *et al.* defined Wave 1 as occurring at around 1 hour, and containing protein cluster 1, 2, 4, and 7 events. One key difference between the phases defined in this work, and the waves defined by Yang *et al.* is that, in the latter, the timing of activation and repression events was set to the beginning of abundance changes, whereas by default Minardo-Model assigns events to when abundance changes are ~50% complete. This highlights the point that a user of Minardo-Model may need to adjust the threshold used to assign events, depending on their analysis goals - for this reason, we provide the code as open source and encourage others to implement different thresholding strategies. Nonetheless, using the 50% threshold, Minardo-Model, similar to Yang *et al.*, identified protein cluster 1 activation to be the earliest one, and classed this event into phase 1. Grouping of protein cluster 4 and 7 events in later phases (phase 2, and early phase 3) is supported because of the observation of transcriptional regulation of key proteins in these clusters, at earlier or simultaneous times (such as Nanog and Stat3, as detailed above).

Wave 2 was defined by Yang *et al.* to occur from 6-24 hours, and contained events from clusters 3, 5, 6, and 8-10. Wave 3 was defined to occur at 36 hours, and contained events from cluster 11 and 12. In our analysis, events from cluster 6 translation activation were in phase 3.

Similar to Yang *et al.*, we found cluster 11 and 12 events to be one of the last translation events, and these events were also distinctly ordered from all other translation events. In summary, the three phases found in the automated ordering generated by Minardo-Model mapped well onto the three waves identified in the manual analysis of Yang *et al.*, demonstrating that our method has potential to help streamline the analysis and interpretation of complex, multiomic data sets.

#### Supplementary references


1. Bezdek, J. C., Ehrlich, R. & Full, W. FCM: The fuzzy c-means clustering algorithm. *Computers & Geosciences* **10,** 191–203 (1984).
2. Humphrey, S. J. *et al.* Dynamic adipocyte phosphoproteome reveals that Akt directly regulates mTORC2. *Cell Metab.* **17,** 1009–1020 (2013).
3. Yang, P. *et al.* Knowledge-Based Analysis for Detecting Key Signaling Events from Time-Series Phosphoproteomics Data. *PLoS Comput. Biol.* **11,** e1004403 (2015).
4. Yang, P. *et al.* Multi-omic Profiling Reveals Dynamics of the Phased Progression of Pluripotency. *Cell Syst.* **8,** 427-445.e10 (2019).
5. Ernst, J. & Bar-Joseph, Z. STEM: a tool for the analysis of short time series gene expression data. *BMC Bioinformatics* **7,** 191 (2006).
6. Ma, D. K. G., Stolte, C., Krycer, J. R., James, D. E. & O'Donoghue, S. I. SnapShot: Insulin/IGF1 Signaling. *Cell* **161,** 948-948.e1 (2015).
7. Kaur, S., Baldi, B., Vuong, J. & O'Donoghue, S. I. A benchmark dataset for analyzing and visualizing the dynamic epiproteome. *Data Brief* **25,** 104000 (2019).
8. Spearman, C. The proof and measurement of association between two things. *Int. J. Epidemiol.* **39,** 1137–1150 (2010).
9. Schulz, M. H. *et al.* DREM 2.0: Improved reconstruction of dynamic regulatory networks from time-series expression data. *BMC Syst. Biol.* **6,** 104 (2012).

10. Ernst, J., Vainas, O., Harbison, C. T., Simon, I. & Bar-Joseph, Z. Reconstructing dynamic regulatory maps. *Mol. Syst. Biol.* **3,** 74 (2007).

11. Watanabe, S. *et al.* Activation of Akt signaling is sufficient to maintain pluripotency in mouse and primate embryonic stem cells. *Oncogene* **25,** 2697–2707 (2006).

12. Altshuler, A. *et al.* RAS Regulates the Transition from Naive to Primed Pluripotent Stem Cells. *Stem Cell Reports* **10,** 1088–1101 (2018).

13. Yu, X. *et al.* Notch signaling activation in human embryonic stem cells is required for embryonic, but not trophoblastic, lineage commitment. *Cell Stem Cell* **2,** 461–471 (2008).

14. Huang, D. W., Sherman, B. T. & Lempicki, R. A. Systematic and integrative analysis of large gene lists using DAVID bioinformatics resources. *Nat. Protoc.* **4,** 44–57 (2009).

15. Zhou, H. *et al.* Conversion of mouse epiblast stem cells to an earlier pluripotency state by small molecules. *J. Biol. Chem.* **285,** 29676–29680 (2010).

16. Johnson, B. V., Shindo, N., Rathjen, P. D., Rathjen, J. & Keough, R. A. Understanding pluripotency--how embryonic stem cells keep their options open. *Mol. Hum. Reprod.* **14,** 513–520 (2008).

17. Kalkan, T. *et al.* Tracking the embryonic stem cell transition from ground state pluripotency. *Development* **144,** 1221–1234 (2017).

18. Simeone, A. & Acampora, D. The role of Otx2 in organizing the anterior patterning in mouse. *Int. J. Dev. Biol.* **45,** 337–345 (2001).
